# Phospholamban and sarcolipin share similar transmembrane zipper motifs that control self-association affinity and oligomer stoichiometry

**DOI:** 10.1101/701797

**Authors:** Nicholas R. DesLauriers, Bengt Svensson, David D. Thomas, Joseph M. Autry

**Author notes:** To whom correspondence should be addressed: Joseph M. Autry, University of Minnesota, Biophysical Technology Center, 6-155 Jackson Hall, 321 Church Street S.E., Minneapolis, MN 55455, USA.

## Abstract

We have characterized the structural determinants of phospholamban (PLB) and sarcolipin (SLN) self-association using site-directed mutagenesis, SDS-PAGE, and fluorescence resonance energy transfer (FRET) microscopy. PLB and SLN are single-pass transmembrane (TM) peptides that are critically involved in regulation of contractility in cardiac and skeletal muscle via reversible inhibition of calcium (Ca) transport by SERCA. PLB and SLN also exhibit ion channel activity *in vitro*, yet the physiological significance of these functions is unknown. Here we have determined that structural insights offered by the tetrameric PLB Cys41 to Leu (C41L) mutation, a mutant with four possible leucine/isoleucine zipper interactions for stabilizing PLB tetramers. Using scanning alanine mutagenesis and SDS-PAGE, we have determined the C41L-PLB tetramer is destabilized by mutation of Leu37 to Ala (L37A) or Ile40 to Ala (I40A), which are the same *a*- and *d*-arm residues stabilizing the PLB pentamer via leucine/isoleucine zippers, highlighting the importance of these two zippers in PLB higher-order oligomerization. The new possible zipper arm in C41L-PLB (N34, C41L, I48) did not contribute to tetramerization. On the other hand, we determined that tetramer conversion back to pentamer was induced by alanine mutation of Ile48, a residue located on the *e*-arm below C41L, implicating steric interaction and restriction are the stabilizing and destabilizing forces that control the distribution between pentamer and tetramer populations. We propose that the *e*-arm and hydrophobic residues in the adjacent *b*-arm act as secondary structural motifs that help control the stoichiometry of PLB oligomerization. FRET microscopy and alanine mutagenesis of SLN residues Val14 (V14A) or Leu21 (L21A) decreased the binding affinity of the SLN‒SLN complex, demonstrating the importance of each residue in mediating self-association. Helical wheel analysis supports a heptad-repeat TM zipper mechanism of SLN oligomerization, similar to the 3.5 residue/turn Leu and Ile zippers found in PLB pentamers. Collectively, our studies add new insights on the conservation of homologous hydrophobic 3-4 pattern of residues in zipper motifs that mediate PLB and SLN self-assembly. We propose that the importance of these apolar, steric interactions in the TM domain are widespread in stabilizing higher-order oligomerization of membrane proteins.

## Introduction

Sarcolipin (SLN)^1^ and phospholamban (PLB) are two key proteins that regulate muscle and cardiac contractility. Both SLN and PLB monomers have a primary regulatory function of inhibiting Ca (Ca) transport of the sarcoplasmic reticulum Ca-transporting ATPase (SERCA). SLN inhibits SERCA in fast-twitch skeletal and atrial muscle, while PLB inhibits SERCA in cardiac and slow-twitch muscle. Adrenaline stimulation of these muscle results in phosphorylation of SLN and PLB, thereby reversing SERCA inhibition and stimulating contractility, with ionotropic and lusitropic effects.

While the primary function of monomeric SLN and PLB is known, both proteins are also capable of self-assembling into higher order oligomers. PLB avidly forms pentamers, and does so in dynamic equilibrium with the monomeric population that actively inhibits SERCA (Wegener and Jones 1984, Simmerman, Kobayashi et al. 1996, Autry and Jones 1997, Cornea, Jones et al. 1997, Kimura, Kurzydlowski et al. 1997, Autry and Jones 1998, Cornea, Autry et al. 2000) Similarly, SLN forms oligomers in detergent micelles, live cell ER (endoplasmic reticulum) membranes, and reconstituted proteoliposomes and tubular lipid crystals (Hellstern, Pegoraro et al. 2001, Becucci, Guidelli et al. 2009, Autry, Rubin et al. 2011, Cao, Wu et al. 2017).

The physiological function of oligomeric SLN and PLB remain unknown. It has proposed that higherorder oligomers act as a conglomerated pool of non-inhibitory PLB or SLN that exist in equilibrium with monomers, which in the non-phosphorylated form actively inhibit SERCA (Autry, Rubin et al. 2011, Shaikh, Sahoo et al. 2016), though the effect of phosphorylation on oligomerization remains unclear. There is also a growing body of evidence that SLN and PLB oligomers possess channel activity. The idea of this second function was proposed when purified PLB was added to lipid bilayers (Kovacs, Nelson et al. 1988, Arkin, Adams et al. 1994, Arkin, Rothman et al. 1995, Oxenoid and Chou 2005, Becucci, Cembran et al. 2009, Smeazzetto, Saponaro et al. 2013, Smeazzetto, Sacconi et al. 2014, Smeazzetto, Tadini-Buoninsegni et al. 2016). PLB was reported to form channels through the bilayers, and the bilayers were reported to be selective for cations (Ca or K^+^). Thus, it was proposed that either (i) PLB acts as a Ca leak channel, acting to inhibit Ca reuptake by SERCA in the sarcoplasmic reticulum (SR), (ii) adding to Ca release from the SR, and/or (iii) providing K^+^ counter-cation flux to help neutralize the large electro-gradient generated during Ca release. PLB expression in *E. coli* has also been reported to decrease cell viability, presumably due to formation of pentameric Ca leak channels (Cook, Huggins et al. 1989). SLN too shows ion channel activity. When reconstituted in lipid bilayers, SLN increases conductivity of small inorganic anions such as chloride (Cl^−^), phosphate (P_i_), and sulfate, and was proposed to serve as a voltage-gated P_i_ channel (Becucci, Cembran et al. 2009, Becucci, Guidelli et al. 2009).

Structural investigation of the mechanics of PLB and SLN oligomerization has helped further elucidate function. It has been demonstrated that one side of the PLB transmembrane (TM) domain contains hydrophobic residues that form zippers in pentamer formation (Simmerman, Kobayashi et al. 1996, Kimura, Kurzydlowski et al. 1997, Adams, Lee et al. 1998). Previous homology modelling has suggested that like its cousin protein, SLN forms higher-order oligomers using leucine/isoleucine zippers. Helical wheel analysis and sequence comparison have indicated that position I17-SLN is analogous to the Ile40PLB zipper position, and indeed when Ile17 was mutated to Ala (I17A), the formation of higher-order oligomers and self-association binding affinity were decreased (Autry, Rubin et al. 2011). Wild-type (WT) SLN expression in the *E. coli* DH5α strain also reduces cell viability, as what had been previously observed with PLB expression (Autry, Rubin et al. 2011). However, expression of the I17A mutant restores bacterial cell viability, suggesting that the I17A mutation disrupts a leucine-isoleucine zipper that stabilizes a toxic SLN ion channel in *E. coli* (Autry, Rubin et al. 2011). Residue-residue interactions in molecular dynamic (MD) simulations have also predicted leucine-isoleucine zippers in SLN dimerization and pentamerization (Cao, Wu et al. 2016, Cao, Wu et al. 2016, Cao, Wu et al. 2017).

In recent studies, homology modelling and MD stimulations have been used to study a model of pentameric SLN channel in membrane. Potential mean force (PMF) energy barrier values indicated permeability of hydrated sodium (Na^+^) and Cl^−^ ions through pentameric SLN. As the narrowest portion of the pore and highest energy barrier for either Na^+^ or Cl^−^ passage correlated with the hydrophobic L21 position, it was hypothesized this residue serves as the ion gate and that hydration expanded the pore size sufficiently enough to allow ion passage—a “hydrophobic gate” (Cao, Wu et al. 2016, Cao, Wu et al. 2017). MD simulations also predicted that phosphorylated SLN pentamers (phospho-Thr5) have greater ion permeability than non-phosphorylated SLN pentamers (Cao, Wu et al. 2016).

Much remains to be understood of the structural motifs for PLB and SLN oligomerization and the functional characteristics of channel activity. Our FRET and SDS-PAGE results help define the TM residue interactions and zipper mechanisms involved in SLN and PLB self-assembly. An improved understanding of how PLB and SLN oligomers form and what regulates their assembly is crucial in determining their physiological function.

## Materials and Methods

### Materials

Chemicals were purchased from Sigma-Aldrich Corp. cDNAs encoding cyan fluorescent protein (CFP) and yellow fluorescent protein (YFP) were purchased from Clontech Com. DNA mutagenesis QuikChange kit was purchased from Stratagene Com. The baculovirus transfer plasmid pAcSG2 was purchased from Orbigen, Inc. BaculoGold baculovirus DNA was purchased from Pharmingen, Com. *Spodoptera frugiperda* (Sf21) insect cells were purchased from Invitrogen. Bacterial expression plasmid pProEx HTc was purchased from Invitrogen. Laemmli-type, pre-cast SDS-PAGE gels were purchased from Bio-Rad Laboratories, Inc. Anti-PLB mouse monoclonal antibody (mAb) 2D12 was purchased afrom Affinity Bioreagents. Anti-tetra-His-tag mouse monoclonal antibody (mAb) IgG1 was purchased from Qiagen Com. (catalog # 34670: 0.2 mg/mL) and used at 1:1000 dilution. The membrane was blocked with 30 mg/mL bovine serum albumin (BSA) in Tris-buffered saline (TBS) at pH 7.4 for 1 hr.

### cDNA cloning, plasmid construction, and site-directed mutagenesis

SLN cDNA was cloned from rabbit fast-twitch muscle mRNA using reverse transcription-polymerase chain reaction (RT-PCR) (Autry, Rubin et al. 2011), finding a cDNA sequence that was the same as previously described (Odermatt, Taschner et al. 1997). cDNA encoding CFP or YFP was then fused to the N-terminus of SLN cDNA using a restriction enzyme/ligation strategy to generate CFP-SLN and YFPSLN cDNA (Winters, Autry et al. 2008, Autry, Rubin et al. 2011). To eliminate dimerization of CFP or YFP, residue Ala206 was mutated to Lys (A206K) (Zacharias, Violin et al. 2002). Alanine mutagenesis of SLN TM residues Val14 or Leu21 was used to create CFP-V14A-SLN or YFP-L21A-SLN.

### Oligomerization assay of C41L-PLN tetramer mutants expressed in Sf21 insect cells

PLB cDNA was cloned from dog left ventricle using RT-PCR (Autry, Rubin et al. 2011), with a sequence the same as previously reported (Fujii, Ueno et al. 1987). C41L mutagenesis was conducted to generate a PLB sequence that forms predominantly tetramers on SDS-PAGE (Simmerman, Kobayashi et al. 1996). Subsequent alanine scanning mutagenesis of TM domain residues L37, I38, L39, I40, L42, and I48 to examine effect on oligomerization. The cDNA of the C41L-PLB mutant and each of its alanine TM mutants were then expressed in Sf21 insect cells via baculovirus infection. SDS-PAGE was used to analyze oligomerization of WT-PLB, C41L-PLB, and TM mutants of C41L-PLB, as expressed in Sf21 cells. Prior to electrophoresis, gel samples were solubilized in 1.0% SDS for 5 min at 23°C. Pre-cast 4–15% acrylamide Laemmli gels were run per manufacturer instructions (BioRad). Immunoblotting was performed using anti-dog PLB mAb 2D12 and secondary antibody with chemiluminescence detection.

### FRET assay of SLN fluorescent fusion proteins expressed in Sf21 insect cells

CFP-SLN, YFP-SLN, CFP-V14A-SLN, and YFP-L21A-SLN proteins were expressed in Sf21 insect cells via recombinant baculovirus (Autry and Jones 1997, Autry and Jones 1998, Autry, Rubin et al. 2011). For FRET microscopy, 6.0 × 10^5^ cells were infected in 60 mm^2^ glass-bottom dishes with a multiplicity of infection (MOI) of 2 viruses/cell for the donor construct (CFP-based fusion protein) and 3 viruses/cell for the acceptor construct (YFP-based fusion protein). FRET was measured 48 hr postinfection, as described below (acceptor photobleaching and spectral unmixing).

FRET microscopy was used to assess interaction between CFP- and YFP-labeled SLN constructs. The epifluorescence microscope, photobleaching protocol, and fluorescence quantification of CFP and YFP fusion proteins were the same as previously described (Autry and Jones 1997, Autry and Jones 1998, Autry, Rubin et al. 2011). The Förster equation was used to calculate energy transfer efficiency:

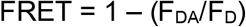

where F_D_ is the fluorescence of the CFP donor in the absence of the YFP acceptor, and F_DA_ is the fluorescence of the CFP donor in the presence of the YFP acceptor. Mean FRET values are reported with ± standard error of the mean (S.E.).

## Results

### Mutagenesis and SDS-PAGE identify zipper and cleft residues that convert C41L-PLB to a tetramer

We expressed PLB in Sf21 insect cells and conducted scanning alanine mutagenesis of the C41L-PLB tetramer to further characterize the role of hydrophobic TM residues in PLB oligomerization. Previous studies have demonstrated the importance of leucine and isoleucine zipper-forming residues Leu37, Ile40, and Leu44 in PLB pentameric stabilization (*a*- and *d*-arms). Here, we identify the structural motif stabilizing the tetramer population and reveal two additional zipper arms involved in oligomerization.

WT-PLB expression in cardiac SR results in a mix of oligomers including dimers, tetramers, and pentamers, as seen on SDS-PAGE (**Figure 1A**, **lane 1**). However, C41L expression hinders pentameric formation and results in predominantly tetrameric population (**Figure 1A**, **lane 2**). Alanine mutations of Leu37 and Ile40 in C41L-PLB subsequently result in absence of the tetramer (**Figure 1**, **lanes 3 and 6**), highlighting the importance of these residues in tetrameric stabilization and indicating they likely play the same role as *a*- and *d*-arm hydrophobic zippers as in PLB pentamers (**Figure 1**). Alanine mutation of residues on the opposite face of the TM helix Ile38, Leu39, and Leu42, meanwhile does not disrupt tetrameric formation (**Figure 1A**, **lanes 4 and 5**).

**Figure 1.**
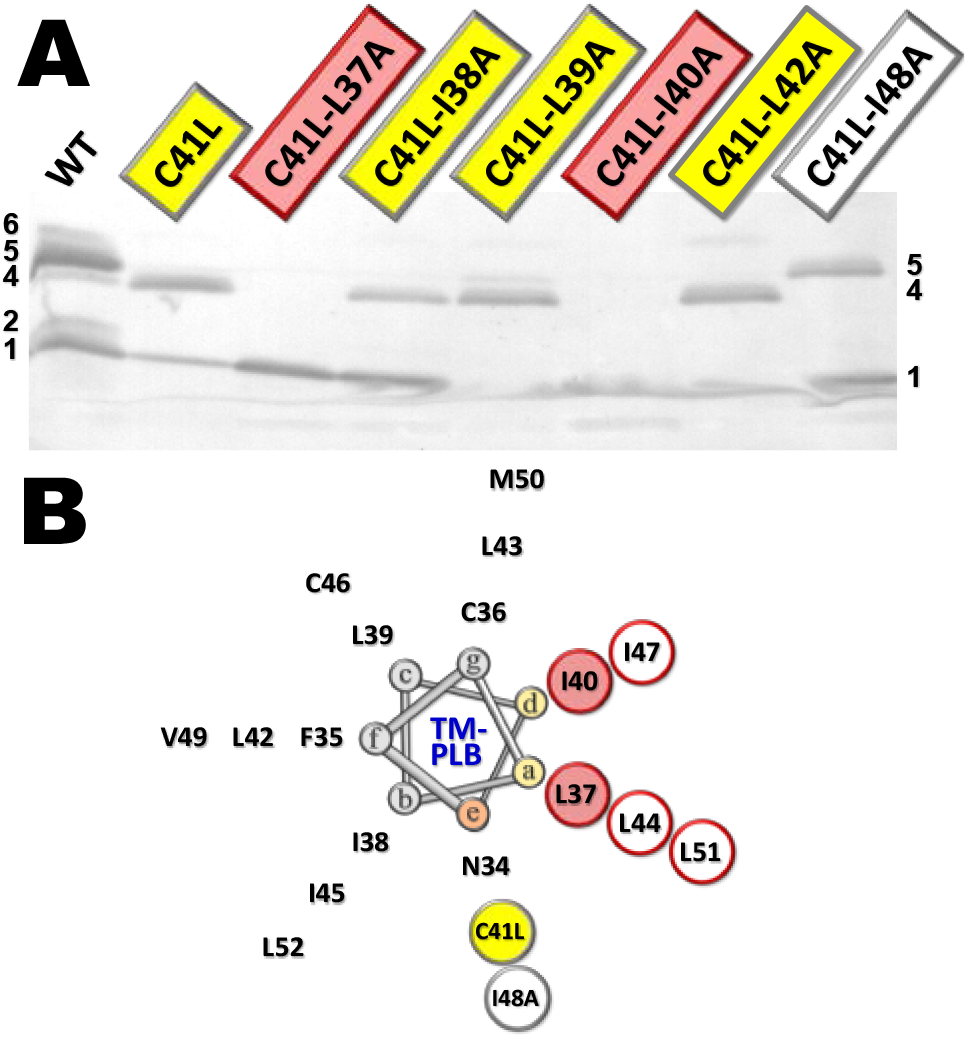
Alanine scanning mutagenesis of tetrameric C41L-PLB. Cys41 was mutated to Leu (C41L) to probe oligomeric interactions of PLB using alanine mutagenesis of residues 37–40, 42, and 48. (A) Immunoblot shows WT-PLB as a mix of oligomeric states, including monomer, dimer, tetramer, pentamer, and higher-order oligomer (probably hexamer). Ala mutants of C41L-PLB that are monomeric are highlighted in *red*, Ala mutants of C14L that are tetrameric are highlighted in *yellow*, and the Ala mutant of C41L-PLB that is pentameric *grey* border. (B) Transmembrane domain of PLB (residues 34–52) as a 3.5 residue-per-turn helix (Simmerman, Kobayashi et al. 1996). Isoleucine and leucine zipper residues are shown in *red*. Ala mutagenesis indicates that the C41L-PLB oligomer is stabilized primarily by *a*- and *d*-arms, and that the *e*-arm determines tetramer versus pentamer oligomeric states.

In addition, we observe that alanine mutation of Ile48 in C41L-PLB diminishes the tetramer population and restores pentamers (**Figure 1**, **lane 8**). This is significant given the position of Ile48 on the *e*-arm below C41L in the proposed helical wheel model (**Figure 1B**). We propose that introduction of the C41L mutation, via conversion from a polar to apolar residue, provides increased hydrophobicity to an additional zipper arm (*e*-arm with C41L and Ile48) which in addition to the native *b*-arm (Ile38, Ile45, Leu52) and *a*- and *d*-arms, totals four zipper arms to stabilize tetramers (**Figure 1B**). While C41L may add hydrophobicity to strengthen *e*-arm zipper interactions, the added bulk of the leucine residue likely provides a sterically counteractive force that destabilizes pentamers. This is supported by the I48A mutation below C41L likely relieving this steric constraint and thus allowing “native” hydrophobic zipper interactions—which we propose to include the same four arms as the C41L tetramer—to again favor pentamer formation. Observation of these changes provide added support for the helical wheel model, the proposed leucine/isoleucine zippers motifs (**Figure 2**), and the importance of hydrophobic van der Waals forces in higher-order oligomer stabilization.

**Figure 2.**
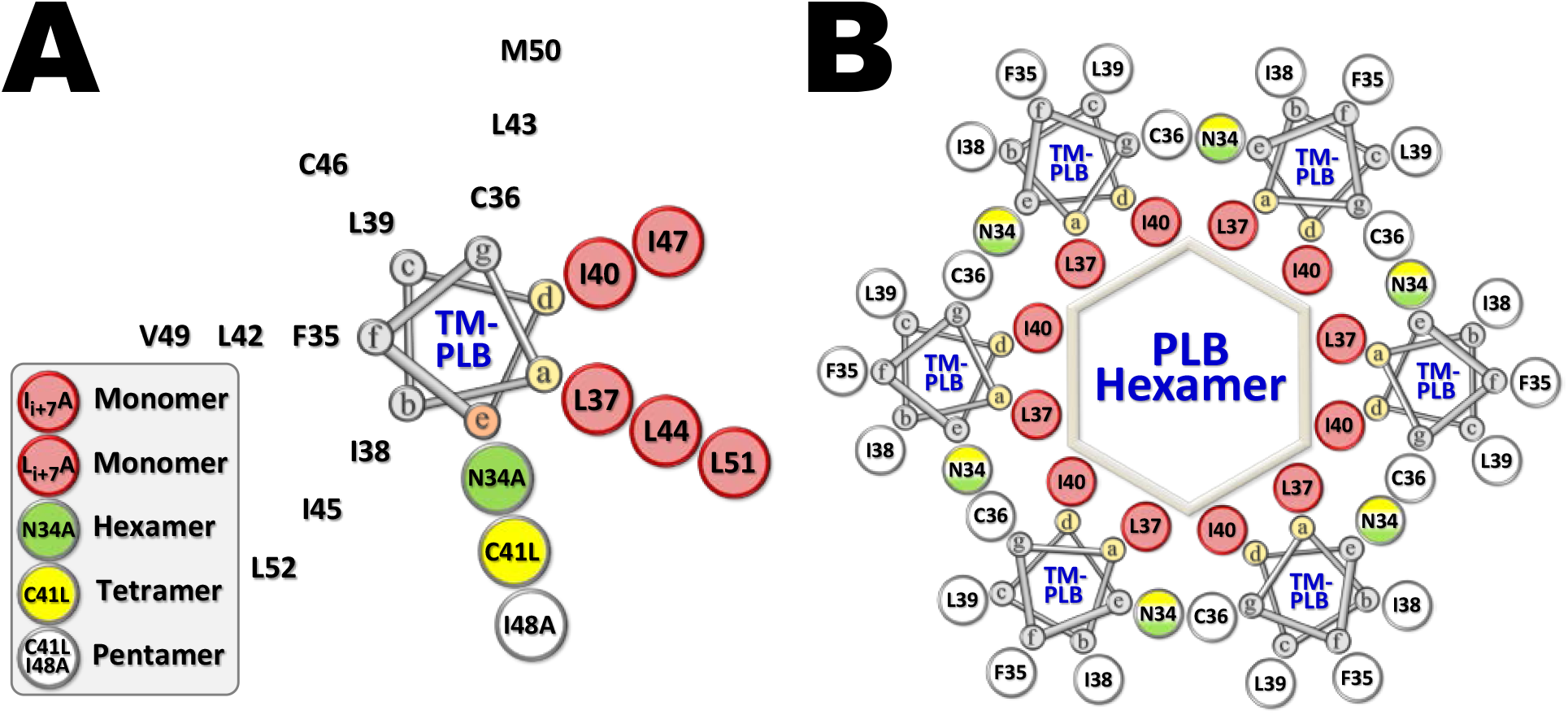
Helical wheel representation of the transmembrane domain of PLB and a structural model for the assembly of the PLB hexamer. (A) Residues 34‒52 of PLB are modeled as a 3.5 residue-per-turn helix with Ile and Leu zipper residues shown in *red*. (Simmerman, Kobayashi et al. 1996). Cys41 was mutated to Leu (C41L) to probe structural determinants of the PLB tetramer. The N34A mutant of PLB shows a mix of pentamers and higher-order oligomers (probably hexamers) on SDS-PAGE (Simmerman, Kobayashi et al. 1996, Kimura, Kurzydlowski et al. 1997). (B) PLB hexamer.

### Mutagenesis and FRET microscopy identify zipper residues that mediate SLN self-association

In previous experiments, the baculovirus system was used to express SLN as a fusion protein with green fluorescent protein derivatives tagged at the N-terminus—CFP-SLN and YFP-SLN (Autry et al 2011). As reported, confocal fluorescence microscopy indicated CFP-SLN and YFP-SLN are localized to the ER when expressed in Sf21 insect cells, and fluorescence microscopy with acceptor-photobleaching indicated a high degree of association between SLN molecules, i.e. that SLN self-associates in live cell membranes. Corroborating this data in a different method of measurement, fluorescence spectroscopy of cell homogenates with detergent dissociation used to disrupt SLN-SLN interactions in isolated Sf21 microsomes demonstrated similar FRET efficiencies.

In this experiment, SLN mutants with alanine substitution to Val14 and to Leu21 were similarly fused with green fluorescent protein derivatives tagged at the N-terminus (YFP-L21A and CFP-V14A), expressed in Sf21 cells, and FRET microscopy conducted on the interactions between each of these mutants with SLN following acceptor-photobleaching. Mean FRET was calculated and binding curves generated using FRET vs acceptor expression (**Figure 3A**). The binding curve for interactions between CFP-SLN and YFP-SLN from the previous experiment is displayed for comparison. Mean FRET between CFP-V14A and YFP-SLN was 0.254 ± 0.009, a decrease of 0.084 ± 0.010 from that between CFP-SLN and YFP-SLN, and K_d_ was 19.7 ± 1.2, a 2-fold reduction in binding affinity from a K_d_ of 8.07 ± 0.31 between CFP-SLN and YFP-SLN. This significant reduction in binding affinity indicates Val14 provides important contribution to SLN self-binding affinity, and the decreased mean FRET that occurs with disruption of this residue alone indicates SLN oligomerization may to some extent be dependent upon it. Mean FRET for interactions between CFP-SLN and YFP-L21A was not different from interactions between CFP-SLN and YFP-SLN, but K_d_ was 13.7 ± 0.8, thus also demonstrating a reduction in binding affinity. Therefore, the L21 residue also plays an important role in stabilization of oligomer formation.

**Figure 3.**
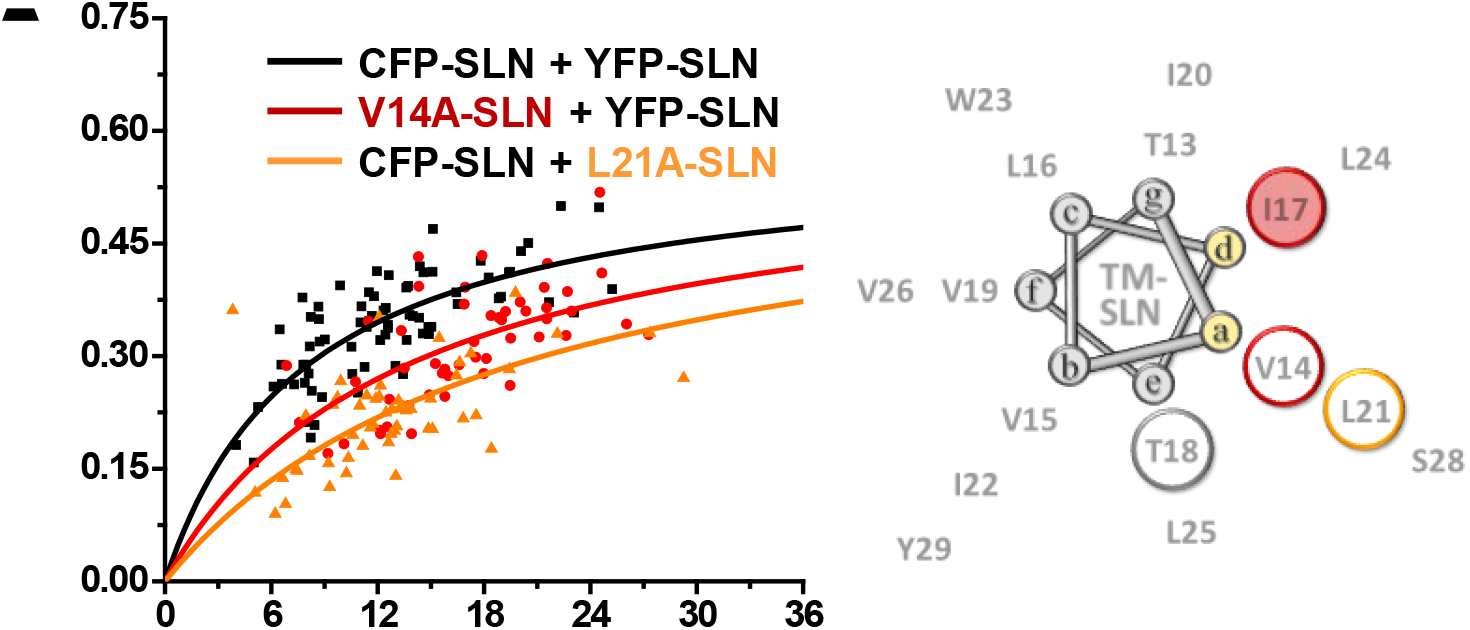
Alanine mutagenesis and FRET analysis of the TM domain of SLN. (A) FRET binding curve. (B) Results are mapped onto the TM domain of SLN modeled as 3.5 residue/turn helix. The T18A mutant (black circle) on the *e*-arm (cleft position) is a loss-of-channel-activity in planar lipid bilayer (Becucci, Guidelli et al. 2009).

FRET analysis from our previous experiment demonstrated that mutation of the Ile17 residue, mapped on the *d*-arm of a proposed 3.5 residue/turn helix model, similarly impairs FRET efficiency. Adding V14 and L21 to this helical wheel analysis places them in the *a*-arm of this model (**Figure 3B**). This adds to a growing body of hydrophobic residues identified as important in self-assembly, and which we propose are stabilizing oligomers via leucine/isoleucine zippers on these two adjacent helical arms.

## Discussion

### Novel structural insights into SLN and PLB self-assembly

In the present study, we investigated interactions of SLN and PLB, using a combination of cellular and biochemical assays. Our results have provided several new insights into the structure and function of these integral membrane proteins.

Results obtained using two heterologous in vivo systems—baculovirus and *E. coli*—indicate that SLN assembles into oligomeric complexes like PLB. In addition to the I17 residue previously identified, our FRET data has identified two additional hydrophobic TM residues, V14 and L21, which are important in SLN higher-order oligomerization (**Figure 3A**). Placing these results in helical wheel analysis supports a leucine/isoleucine zipper mechanism of oligomerization, with at least two zippers located on the *a*- and *d*-arms of a repeating heptad coiled-coil helix (**Figure 3B**). These proposed SLN zippers bear striking similarity to the two leucine/isoleucine zippers on the *a*- and *d*-arms of pentameric PLB, previously identified via disruption of PLB pentamer on gel from mutation of L37, I40, L44, I47, L51 (Simmerman, Kobayashi et al. 1996). Our SDS-PAGE gel results here have also identified *a*- and *d*-arm residues of a PLB tetramer as necessary for stabilization as well as two other potential zippers on the *c*- and *e*-arms (**Figure 1**, **Figure 2**). Collectively, these results provide strong evidence that cousin proteins SLN and PLB share a conserved motif of hydrophobic leucine/isoleucine zippers stabilizing higher-order oligomers.

The discovery here of a “steric switch” capable of converting between PLB tetramer and pentamer, and an examination of the mechanics of how this switch impacts oligomeric order, further supports a hydrophobic zipper mechanism of oligomerization centered on *a*- and *d*-arm interactions. Mutation of *a*- and *d*-arm residues have the greatest evidence of disrupting higher-order oligomerization (Error! Reference source not found.**B**, Error! Reference source not found.**B**) and thus may be the point of strongest contact between monomeric units in higher-order oligomer formation. However, with an increasing oligomer number—i.e., tetramer to pentamer—an increasing angle between monomeric units would require a shift in monomeric contact toward the *e*-arm. Accordingly, the steric bulk introduced at C41L on the *e*-arm likely provides a barrier to any expanding angle between monomers and restricts C41L-PLB to a tetramer, with the tetramer further strengthened by zippers between the *e*-arm (mutant) and *b*-arm (native). Interestingly, a hexameric form of PLN has been observed with N34A mutation—a residue also lying on the *e*-arm (Simmerman, Kobayashi et al. 1996). With reduction of steric bulk on the *e*-arm from this mutation, hydrophobic zipper interactions may be able to maintain a greater angle between monomers, resulting in the preponderance of hexamers rather than pentamers. Taken together, these examples demonstrate the *e*-arm of PLB as an important “steric switch” position and corroborates the hydrophobic zipper model of oligomerization (**Figure 1**, **Figure 2**, **Figure 4**).

**Figure 4.**
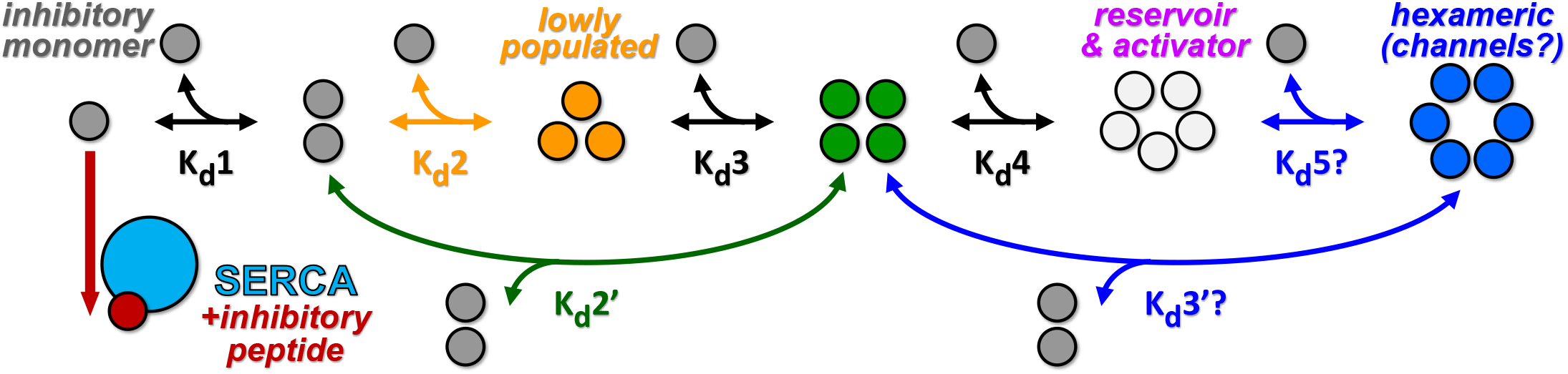
Summary schematic for oligomeric interactions of SERCA and regulatory peptides PLB or SLN.

Notably, previous experiments which truncated the polar upper-TM region of an engineered, watersoluble form of PLB found that this also served as a switch from pentamer to tetramer, which could suggest a greater role for polar forces in higher-order PLB oligomerization (Slovic, Lear et al. 2005). However, emerging research has also continued to demonstrate the prominent role of van der Waal forces in stabilizing peptide oligomers and an engineered PLB-like molecule devoid of strongly polar residues has even been shown to easily stabilize as a pentamer (Mravic, Thomaston et al. 2019), which we suggest is secondary to hydrophobic zipper interactions. The evidence presented here show PLB and SLN as examples supporting the prominent role of hydrophobic interactions in small peptide oligomerization as well as determination of oligomer order.

### Proposed hypothesis on PLB and SLN pore size and channel activity

With SLN known to increase conductance in membranes (Becucci, Cembran et al. 2009, Becucci, Guidelli et al. 2009), and with recent molecular dynamic simulations demonstrating the SLN pentamer as capable of small ion transport (Cao, Wu et al. 2016, Cao, Wu et al. 2017), this adds to a growing body of evidence of SLN oligomers having ion channel capacity. Given an *anionic* conductance in membranes, we predict that SLN forms P_i_ or Cl^−^ channels. A modest population of SLN P_i_ channels, especially upon phosphorylation, may be physiologic in design to allow for a controlled amount of P_i_ import and Ca precipitation in muscle SR. Cl^−^ currents have also been measured in the SR (Tanifuji, Sokabe et al. 1987, Rousseau 1989), though no Cl^−^ channel yet identified. Interestingly, an anti-PLB antibody has been observed to halt Cl^−^ current across vesicles from the SR (Decrouy, Juteau et al. 1996), though it is possible the antibody was cross-reacting with SLN channels.

### Do PLB or SLN act as a hexameric or higher-order oligomeric ion channel?

While there is growing evidence that PLB can acts as a cation channel, PLB pentamers are predicted to have too narrow of a pore to conduct hydrated Ca (Arkin, Rothman et al. 1995). We have demonstrated that WT-PLB can form hexamers in on SDS-PAGE (**Figure 1A**, **lane 1**), along with additional higherorder oligomers (**Supplementary Figure S1**). Given that current structural models indicated that PLB pentamers are likely too narrow to form channels for hydrated Ca ions; we predict that higher-order oligomers of PLB, such as hexamers or larger, are capable of pore formation (**Figure 2**, **Figure 4**). As noted previously, the N34A mutant of PLB also forms a high-order oligomer on SDS-PAGE (probably hexamer), with alanine mutation at this cleft position (*e*-arm) facilitating formation by alleviating steric hindrance (Simmerman, Kobayashi et al. 1996). Ultracentrifugation and cross-linking have also detected PLB hexamers and decamers in detergent (Vorherr, Wrzosek et al. 1993, Hellstern, Pegoraro et al. 2001), while FRET has detected a significant fraction of higher-order PLB oligomers with 10‒12 subunits when reconstituted in dioleoylphosphatidylcholine (DOPC) bilayers (Reddy, Jones et al. 1999). Much focus and study has been on the PLB pentamer given its preponderance in SDS-PAGE assays. However, it is possible that the presence of SDS does not favor formation of high-order oligomers of PLB. we demonstrate greater temperature alone has the capacity to reduce the hexamer population to undetectable, while the pentameric population remains (**Supplementary Figure S1**). Thus, PLB hexamers and species with a greater number of subunits may well exist SR membranes in greater populaiton than previously appreciated.

There is an increasing number of protein channels built from TM helices proving to demonstrate a fluid rather than set oligomerization number in membranes, with some proposed capable of spanning up to a dodecameric order (Saravanan and Bhattacharjya 2011, Bortolus, De Zotti et al. 2013, Serban, Breen et al. 2018). Similarly, hexameric and a spectrum of greater order PLB channel-forming oligomers may well be in an equilibrium with lower-order oligomers, resulting in a modest population of PLB Ca channels that may be physiologic in design to allow for controlled flux of Ca or K ions across the SR membrane. Given the demonstrated range of PLB higher-order oligomers and the facility with which steric alterations can tune zipper interactions and control oligomer order (**Figure 1A**), this fluid rather than static model of channel oligomer model is conceivable.

Given homologous structure and function, we hypothesize SLN is also capable of forming hexameric and higher-order oligomeric channels. SLN has been shown to be capable of hexamer formation from ultracentrifugation (Hellstern, Pegoraro et al. 2001), though further evidence is needed on the capacity of SLN for higher-order oligomerization. Though we favor a hexameric or greater oligomer number for SLN acting as a P_i_ selective-channel, yet a Cl^−^ selective-channel of SLN with a lower stoichiometry is also conceivable given the smaller anion size.

### Role of PLB pentamer

If hexamer and greater order PLB oligomers are required to form calcium channels, what is the physiologic role of pentameric PLB; the oligomeric species with the greatest preponderance on gels and most studied? There are number of studies providing evidence that PLB pentamers regulate SERCA, including a crystal structure of SERCA interaction with PLB pentamer and a demonstration that SERCA Vmax is increased with a higher concentration of PLN pentamer (Reddy, Cornea et al. 2003, Trieber, Douglas et al. 2005, Stokes, Pomfret et al. 2006, Glaves, Primeau et al. 2019). The Young group proposes that the PLN pentamer sits at the calcium entry funnel of SERCA when bound and its basic residues may thus perturb the surrounding lipid bilayer, resulting in greater mobility of SERCA catalytic pumps and an increased turnover capacity (Glaves, Primeau et al. 2019). Evidence suggesting functions of the PLB pentamer beyond direct regulation of SERCA have also been brought forth, such as its capacity to attenuate PKA to delay the phosphorylation of the PLB monomer (Wittmann, Lohse et al. 2015).

Interestingly, there is also a growing body of evidence on the presence of “chansporters”— complexes between two structurally unrelated but interacting channel and transporter proteins—throughout various different physiologic systems in the body. The neuronal K_v_7.2/7.3 potassium channel, for example, has been found capable of interacting with two different sodium-coupled neurotransmitter transporters, where the potassium channel may be enhancing sodium transport while reducing the depolarization effect from sodium influx (Bartolomé-Martín, Ibáñez et al. 2019). In this context and with the evidence above, it becomes quite conceivable that PLB and SLN pentamers may interact with SERCA in chansporter-like complexes in order to modulate Ca influx into the SR.

The standing paradigm of PLB and SLN self-assembly equilibrium has long been that PLB/SLN monomers inhibit SERCA, which can pentamerize to becomes inactive. While oligomerization may in part serve a storage function for monomers, we believe self-assembly equilibrium is likely far more complex, with a fluid range of oligomer order possible including pentamers, hexamers, and greater, and where these oligomer species carry multiple important functions including SERCA regulation and ion channel activity.

### Proposed physiological function of PLB and SLN channels and medical relevance

We propose a physiologic model whereby PLB and SLN oligomeric channels serve as regulatory mechanisms of muscle contraction. Given ion concentrations in the cell, we predict that a Ca-leak PLB channel would inhibit muscle contraction, while a P_i_ uptake SLN channel would activate contraction. A PLB Ca-leak channel would be conceivable as a mechanism to quickly remove Ca to improve cycling during rapid contraction. A mechanism by which SLN P_i_ uptake channels activate muscle contraction or improve contractility is also conceivable. Inorganic P_i_ levels can increase to >30 mM in contracting muscle cells, and this P_i_ enters the SR to form dissolvable Ca-Pi precipitates to assist calsequestrin with Ca storage (Dutka, Cole et al. 2005). High concentrations of inorganic P_i_ also inhibit myosin (Debold, Walcott et al. 2013). P_i_ uptake channels may thus provide a regulated mechanism of increasing P_i_ in the SR lumen during contraction that can help store Ca to increase contractility or maintain contractility in fatiguing muscle, while ATP is preserved from reduced Ca release and inhibition of myosin (Debold, Walcott et al. 2013). P_i_ uptake channels may thus provide a regulated mechanism of increasing P_i_ in the SR during contraction that can help store Ca to increase contractility or maintain contractility in fatiguing muscle, while ATP is preserved from reduced Ca release and inhibition of myosin.

The proposed roles of SLN and PLB channels in Ca cycling and muscle contractility would have important medical relevance and implications in heart disease. Pathologic Ca leakage out of the SR has been shown to lead to atrial fibrillation (Sood, Chelu et al. 2008, Shan, Xie et al. 2012) and inhibition of this Ca leakage shown to correct arrhythmias (Mohamed, Hartmann et al. 2018). Thus a PLB channel serving as a Ca leakage mechanism would have immediate relevance in pathophysiology or therapeutics. It is also known that decreased SLN expression and enhanced SERCA expression are common in atrial fibrillation, likely causing disruption of Ca cycling in and out of the SR (Shanmugam, Molina et al. 2011). Might this be not only because of SLN’s absent inhibition of SERCA, but also because of the absence of SLN P_i_ channels, diminishing the capacity of Ca storage in the SR in fatiguing muscle? Indeed, if PLB and SLN channels are involved in the Ca cycling of the SR, then formation of these channels may be directly involved in the pathophysiology of any number of other aberrant Ca cycling diseases, or if not directly aberrant themselves may hold promise as therapeutic targets to reverse problems with other components of the Ca cycling machinery. It thus becomes imperative to fully elucidate the role of these two important regulatory proteins.

## Summary

In this research, we have used cellular and biochemical assays to contribute new structural insights to PLB and SLN self-oligomerization. Our studies have added to understanding of the hydrophobic TM residues involved in PLB and SLN self-assembly, in particular demonstrating the conservation of leucine/isoleucine zipper motifs between these cousin proteins and emphasizing the importance of hydrophobic interactions in oligomerization. Adding observation of effects of PLB and SLN oligomers in *E. coli* to a growing understanding of structure, known conductance capacities, and molecular simulations, we have proposed that these oligomers can form ion channels that have a physiologic role in muscle contraction regulation. Continued study of the structure and function of PLB and SLN oligomers is needed to better elucidate their physiologic role, as this may have wide-ranging implications in Ca cycling and play an important role in both pathophysiology and potential therapeutics in muscle and cardiac diseases.

## Acknowledgments

We thank Octavian Cornea, Sarah Blakely-Anderson, and Destiny Ziebol for administrative support. Spectrophotometric assays were performed in the Biophysical Technology Center at the University of Minnesota Department of Biochemistry, Molecular Biology, and Biophysics.

## Author contributions

Project design: J.M.A.

Project administration: J.M.A. and D.D.T.

Project funding: D.D.T., J.M.A., J.E.R., N.R.D., L.M.E.-F., and S.J.V.

Manuscript writing: N.R.D., J.M.A., J.E.R., B.S., L.M.E.-F., and S.J.V.

Manuscript figures: J.M.A. and B.S.

DNA mutagenesis: J.M.A., D.L.W., and N.R.D.

Bacterial cell expression: J.M.A. and N.R.D.

Insect cell expression: J.M.A., J.E.R., and D.L.W.

FRET microscopy: J.E.R. and J.M.A.

Molecular modeling: B.S. and L.M.E.-F.

## Footnotes

1 **Abbreviations and Acronyms:** Amp, ampicillin; Ca, calcium; CAT; chloramphenicol acetyltransferase; CFP, cyan fluorescent protein; Cl^−^, chloride; DOPC, dioleoylphosphatidylcholine; ER, endoplasmic reticulum; FRET, fluorescence resonance energy transfer; his-CAT, chloramphenicol transferase with a 6-residue histidine tag fused to the N-terminus; his-PLB, phospholamban with a 6-residue histidine tag fused to the N-terminus; his-SLN, sarcolipin with a 6-residue histidine tag fused to the N-terminus; IPTG, isopropyl β-D-1-thiogalactopyranoside; LB, lysogeny broth; MD, molecular dynamics; MOI, multiplicity of infection; Na^+^, sodium; PLB, phospholamban; PMF, potential mean force; S.E., standard error of the mean; RT-PCR, reverse transcriptase polymerase chain reaction; SERCA, sarcoplasmic reticulum Ca^2+^-transporting ATPase; Sf21, *Spodoptera frugiperda* insect cells; SLN, sarcolipin; SR, sarcoplasmic reticulum; TM, transmembrane; YFP, yellow fluorescent protein; WT, wild-type.

## References

Adams, P. D., A. S. Lee, A. T. Brunger and D. M. Engelman (1998). “Models for the transmembrane region of the phospholamban pentamer: which is correct?” Ann N Y Acad Sci 853: 178–185.

Arkin, I. T., P. D. Adams, K. R. MacKenzie, M. A. Lemmon, A. T. Brunger and D. M. Engelman (1994). “Structural organization of the pentameric transmembrane alpha-helices of phospholamban, a cardiac ion channel.” EMBO J 13(20): 4757–4764.

Arkin, I. T., M. Rothman, C. F. Ludlam, S. Aimoto, D. M. Engelman, K. J. Rothschild and S. O. Smith (1995). “Structural model of the phospholamban ion channel complex in phospholipid membranes.” J Mol Biol 248(4): 824–834.

Autry, J. M. and L. R. Jones (1997). “Functional co-expression of the canine cardiac Ca2+ pump and phospholamban in Spodoptera frugiperda (Sf21) cells reveals new insights on ATPase regulation.” J Biol Chem 272(25): 15872–15880.

Autry, J. M. and L. R. Jones (1998). “High-level coexpression of the canine cardiac calcium pump and phospholamban in Sf21 insect cells.” Ann N Y Acad Sci 853: 92–102.

Autry, J. M., J. E. Rubin, S. D. Pietrini, D. L. Winters, S. L. Robia and D. D. Thomas (2011). “Oligomeric interactions of sarcolipin and the Ca-ATPase.” J Biol Chem 286(36): 31697–31706.

Bartolomé-Martín, D., I. Ibáñez, D. Piniella, E. Martínez-Blanco, S. G. Pelaz and F. Zafra (2019). “Identification of potassium channel proteins Kv7.2/7.3 as common partners of the dopamine and glutamate transporters DAT and GLT-1.” Neuropharmacology.

Becucci, L., A. Cembran, C. B. Karim, D. D. Thomas, R. Guidelli, J. Gao and G. Veglia (2009). “On the function of pentameric phospholamban: ion channel or storage form?” Biophys J 96(10): L60–62.

Becucci, L., R. Guidelli, C. B. Karim, D. D. Thomas and G. Veglia (2009). “The role of sarcolipin and ATP in the transport of phosphate ion into the sarcoplasmic reticulum.” Biophys J 97(10): 2693–2699.

Bortolus, M., M. De Zotti, F. Formaggio and A. L. Maniero (2013). “Alamethicin in bicelles: orientation, aggregation, and bilayer modification as a function of peptide concentration.” Biochim Biophys Acta 1828(11): 2620–2627.

Cao, Y., X. Wu, I. Lee and X. Wang (2016). “Molecular dynamics of water and monovalent-ions transportation mechanisms of pentameric sarcolipin.” Proteins 84(1): 73–81.

Cao, Y., X. Wu, X. Wang, H. Sun and I. Lee (2016). “Transmembrane dynamics of the Thr-5 phosphorylated sarcolipin pentameric channel.” Arch Biochem Biophys 604: 143–151.

Cao, Y., X. Wu, R. Yang, X. Wang, H. Sun and I. Lee (2017). “Self-assembling study of sarcolipin and its mutants in multiple molecular dynamic simulations.” Proteins 85(6): 1065–1077.

Cook, E. A., J. P. Huggins, G. Sathe, P. J. England and J. R. Piggott (1989). “The expression of canine cardiac phospholamban in heterologous systems.” Biochem J 264(2): 533–538.

Cornea, R. L., J. M. Autry, Z. Chen and L. R. Jones (2000). “Reexamination of the role of the leucine/isoleucine zipper residues of phospholamban in inhibition of the Ca2+ pump of cardiac sarcoplasmic reticulum.” J Biol Chem 275(52): 41487–41494.

Cornea, R. L., L. R. Jones, J. M. Autry and D. D. Thomas (1997). “Mutation and phosphorylation change the oligomeric structure of phospholamban in lipid bilayers.” Biochemistry 36(10): 2960–2967.

Debold, E. P., S. Walcott, M. Woodward and M. A. Turner (2013). “Direct observation of phosphate inhibiting the force-generating capacity of a miniensemble of Myosin molecules.” Biophys J 105(10): 2374–2384.

Decrouy, A., M. Juteau, S. Proteau, J. Teijiera and E. Rousseau (1996). “Biochemical regulation of sarcoplasmic reticulum Cl-channel from human atrial myocytes: involvement of phospholamban.” J Mol Cell Cardiol 28(4): 767–780.

Dutka, T. L., L. Cole and G. D. Lamb (2005). “Calcium phosphate precipitation in the sarcoplasmic reticulum reduces action potential-mediated Ca2+ release in mammalian skeletal muscle.” Am J Physiol Cell Physiol 289(6): C1502–1512.

Fujii, J., A. Ueno, K. Kitano, S. Tanaka, M. Kadoma and M. Tada (1987). “Complete complementary DNA-derived amino acid sequence of canine cardiac phospholamban.” J Clin Invest 79(1): 301–304.

Glaves, J. P., J. O. Primeau, L. M. Espinoza-Fonseca, M. J. Lemieux and H. S. Young (2019). “The phospholamban pentamer alters function of the sarcoplasmic reticulum calcium pump SERCA.” Biophys J 116(4): 633–647.

Hellstern, S., S. Pegoraro, C. B. Karim, A. Lustig, D. D. Thomas, L. Moroder and J. Engel (2001). “Sarcolipin, the shorter homologue of phospholamban, forms oligomeric structures in detergent micelles and in liposomes.” J Biol Chem 276(33): 30845–30852.

Kimura, Y., K. Kurzydlowski, M. Tada and D. H. MacLennan (1997). “Phospholamban inhibitory function is activated by depolymerization.” J Biol Chem 272(24): 15061–15064.

Kovacs, R. J., M. T. Nelson, H. K. Simmerman and L. R. Jones (1988). “Phospholamban forms Ca2+selective channels in lipid bilayers.” J Biol Chem 263(34): 18364–18368.

Mohamed, B. A., N. Hartmann, P. Tirilomis, K. Sekeres, W. Li, S. Neef, C. Richter, E. M. Zeisberg, L. Kattner, M. Didié, K. Guan, J. D. Schmitto, S. E. Lehnart, S. Luther, N. Voigt, T. Seidler, S. Sossalla, G. Hasenfuss and K. Toischer (2018). “Sarcoplasmic reticulum calcium leak contributes to arrhythmia but not to heart failure progression.” Sci Transl Med 10(458).

Mravic, M., J. L. Thomaston, M. Tucker, P. E. Solomon, L. Liu and W. F. DeGrado (2019). “Packing of apolar side chains enables accurate design of highly stable membrane proteins.” Science 363(6434): 1418–1423.

Odermatt, A., P. E. Taschner, S. W. Scherer, B. Beatty, V. K. Khanna, D. R. Cornblath, V. Chaudhry, W. C. Yee, B. Schrank, G. Karpati, M. H. Breuning, N. Knoers and D. H. MacLennan (1997). “Characterization of the gene encoding human sarcolipin (SLN), a proteolipid associated with SERCA1: absence of structural mutations in five patients with Brody disease.” Genomics 45(3): 541–553.

Oxenoid, K. and J. J. Chou (2005). “The structure of phospholamban pentamer reveals a channel-like architecture in membranes.” Proc Natl Acad Sci U S A 102(31): 10870–10875.

Reddy, L. G., R. L. Cornea, D. L. Winters, E. McKenna and D. D. Thomas (2003). “Defining the molecular components of calcium transport regulation in a reconstituted membrane system.” Biochemistry 42(15): 4585–4592.

Reddy, L. G., L. R. Jones and D. D. Thomas (1999). “Depolymerization of phospholamban in the presence of calcium pump: a fluorescence energy transfer study.” Biochemistry 38(13): 3954–3962.

Rousseau, E. (1989). “Single chloride-selective channel from cardiac sarcoplasmic reticulum studied in planar lipid bilayers.” J Membr Biol 110(1): 39–47.

Saravanan, R. and S. Bhattacharjya (2011). “Oligomeric structure of a cathelicidin antimicrobial peptide in dodecylphosphocholine micelle determined by NMR spectroscopy.” Biochim Biophys Acta 1808(1): 369–381.

Serban, A. J., I. L. Breen, H. Q. Bui, M. Levitus and R. M. Wachter (2018). “Assembly-disassembly is coupled to the ATPase cycle of tobacco Rubisco activase.” J Biol Chem 293(50): 19451–19465.

Shaikh, S. A., S. K. Sahoo and M. Periasamy (2016). “Phospholamban and sarcolipin: Are they functionally redundant or distinct regulators of the Sarco(Endo)Plasmic Reticulum Calcium ATPase?” J Mol Cell Cardiol 91: 81–91.

Shan, J., W. Xie, M. Betzenhauser, S. Reiken, B. X. Chen, A. Wronska and A. R. Marks (2012). “Calcium leak through ryanodine receptors leads to atrial fibrillation in 3 mouse models of catecholaminergic polymorphic ventricular tachycardia.” Circ Res 111(6): 708–717.

Shanmugam, M., C. E. Molina, S. Gao, R. Severac-Bastide, R. Fischmeister and G. J. Babu (2011). “Decreased sarcolipin protein expression and enhanced sarco(endo)plasmic reticulum Ca2+ uptake in human atrial fibrillation.” Biochem Biophys Res Commun 410(1): 97–101.

Simmerman, H. K., Y. M. Kobayashi, J. M. Autry and L. R. Jones (1996). “A leucine zipper stabilizes the pentameric membrane domain of phospholamban and forms a coiled-coil pore structure.” J Biol Chem 271(10): 5941–5946.

Simmerman, H. K. B. Y. M. Kobayashi, J. M. Autry and L. R. Jones (1996). “A Leucine Zipper Stabilizes the Pentameric Membrane Domain of Phospholamban and Forms a Coiled-coil Pore Structure.” J. Biol. Chem. 271(10): 5941–5946.

Slovic, A. M., J. D. Lear and W. F. DeGrado (2005). “De novo design of a pentameric coiled-coil: decoding the motif for tetramer versus pentamer formation in water-soluble phospholamban.” J Pept Res 65(3): 312–321.

Smeazzetto, S., A. Sacconi, A. L. Schwan, G. Margheri and F. Tadini-Buoninsegni (2014). “Binding of a monoclonal antibody to the phospholamban cytoplasmic domain interferes with the channel activity of phospholamban reconstituted in a tethered bilayer lipid membrane.” Langmuir 30(34): 10384–10388.

Smeazzetto, S., A. Saponaro, H. S. Young, M. R. Moncelli and G. Thiel (2013). “Structure-function relation of phospholamban: modulation of channel activity as a potential regulator of SERCA activity.” PLoS One 8(1): e52744.

Smeazzetto, S., F. Tadini-Buoninsegni, G. Thiel, D. Berti and C. Montis (2016). “Phospholamban spontaneously reconstitutes into giant unilamellar vesicles where it generates a cation selective channel.” Phys Chem Chem Phys 18(3): 1629–1636.

Sood, S., M. G. Chelu, R. J. van Oort, D. Skapura, M. Santonastasi, D. Dobrev and X. H. Wehrens (2008). “Intracellular calcium leak due to FKBP12.6 deficiency in mice facilitates the inducibility of atrial fibrillation.” Heart Rhythm 5(7): 1047–1054.

Stokes, D. L., A. J. Pomfret, W. J. Rice, J. P. Glaves and H. S. Young (2006). “Interactions between Ca2+ATPase and the pentameric form of phospholamban in two-dimensional co-crystals.” Biophys J 90(11): 4213–4223.

Tanifuji, M., M. Sokabe and M. Kasai (1987). “An anion channel of sarcoplasmic reticulum incorporated into planar lipid bilayers: single-channel behavior and conductance properties.” J Membr Biol 99(2): 103–111.

Trieber, C. A., J. L. Douglas, M. Afara and H. S. Young (2005). “The effects of mutation on the regulatory properties of phospholamban in co-reconstituted membranes.” Biochemistry 44(9): 3289–3297.

Vorherr, T., A. Wrzosek, M. Chiesi and E. Carafoli (1993). “Total synthesis and functional properties of the membrane-intrinsic protein phospholamban.” Protein Sci 2(3): 339–347.

Wegener, A. D. and L. R. Jones (1984). “Phosphorylation-induced mobility shift in phospholamban in sodium dodecyl sulfate-polyacrylamide gels. Evidence for a protein structure consisting of multiple identical phosphorylatable subunits.” J Biol Chem 259(3): 1834–1841.

Winters, D. L., J. M. Autry, B. Svensson and D. D. Thomas (2008). “Interdomain fluorescence resonance energy transfer in SERCA probed by cyan-fluorescent protein fused to the actuator domain.” Biochemistry 47(14): 4246–4256.

Wittmann, T., M. J. Lohse and J. P. Schmitt (2015). “Phospholamban pentamers attenuate PKA-dependent phosphorylation of monomers.” J Mol Cell Cardiol 80: 90–97.

Zacharias, D. A., J. D. Violin, A. C. Newton and R. Y. Tsien (2002). “Partitioning of lipid-modified monomeric GFPs into membrane microdomains of live cells.” Science 296(5569): 913–916.

