## Supplemental Figure S1 for "Phospholamban and sarcolipin share similar transmembrane zipper motifs that control self-association affinity and oligomer stoichiometry"

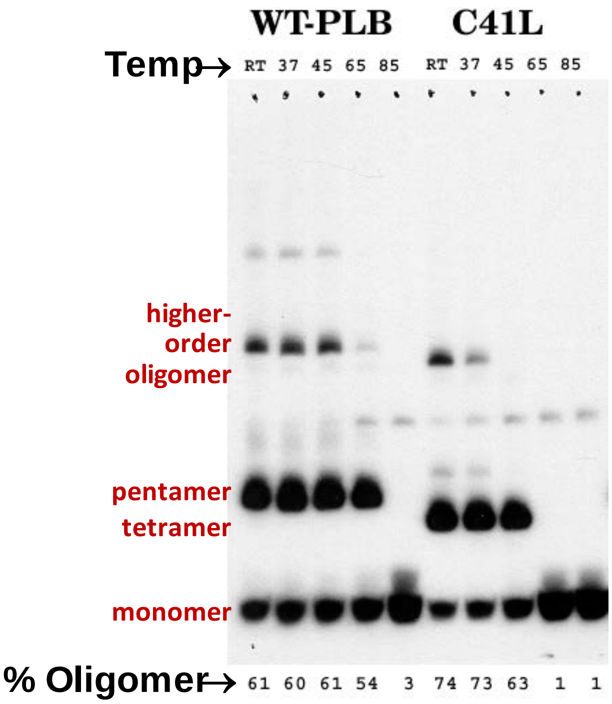


**Figure Supplement S1. Temperature sensitivity of PLB pentamer, tetramer, and higher-order oligomers.**
